# *KAT6A* mutations in Arboleda-Tham syndrome drive epigenetic regulation of posterior *HOXC* cluster

**DOI:** 10.1101/2023.08.03.550595

**Authors:** Meghna Singh, Sarah Spendlove, Angela Wei, Leroy Bondhus, Aileen Nava, Francisca N. de L. Vitorino, Seth Amano, Jacob Lee, Gesenia Echeverria, Dianne Gomez, Benjamin A. Garcia, Valerie A. Arboleda

## Abstract

Arboleda-Tham Syndrome (ARTHS) is a rare genetic disorder caused by heterozygous, *de novo* truncating mutations in *Lysine(K) acetyltransferase 6A* (*KAT6A)*. ARTHS is clinically heterogeneous and characterized by several common features including intellectual disability, developmental and speech delay, hypotonia and affects multiple organ systems. *KAT6A* is highly expressed in early development and plays a key role in cell-type specific differentiation. KAT6A is the enzymatic core of a histone-acetylation protein complex, however the direct histone targets and gene regulatory effects remain unknown. In this study, we use ARTHS patient (n=8) and control (n=14) dermal fibroblasts and perform comprehensive profiling of the epigenome and transcriptome caused by *KAT6A* mutations. We identified differential chromatin accessibility within the promoter or gene body of 23%(14/60) of genes that were differentially expressed between ARTHS and controls. Within fibroblasts, we show a distinct set of genes from the posterior *HOXC* gene cluster (*HOXC10*, *HOXC11*, *HOXC-AS3*, HOXC-AS2, HOTAIR) that are overexpressed in ARTHS and are transcription factors critical for early development body segment patterning. The genomic loci harboring HOXC genes are epigenetically regulated with increased chromatin accessibility, high levels of H3K23ac, and increased gene-body DNA methylation compared to controls, all of which are consistent with transcriptomic overexpression. Finally, we used unbiased proteomic mass spectrometry and identified two new histone post-translational modifications (PTMs) that are disrupted in ARTHS: H2A and H3K56 acetylation. Our multi-omics assays have identified novel histone and gene regulatory roles of *KAT6A* in a large group of ARTHS patients harboring diverse pathogenic mutations. This work provides insight into the role of KAT6A on the epigenomic regulation in somatic cell types.

## Introduction

Arboleda-Tham Syndrome (ARTHS, OMIM#616268) is a rare genetic disorder caused by *de novo* heterozygous mutations in the *lysine(K) acetyltransferase 6A* (*KAT6A*, a.k.a. *MYST3* and *MOZ*) gene. ARTHS is characterized by intellectual disability, developmental and speech delays, hypotonia and congenital heart defects (Arboleda et al. 2015; Tham et al. 2015; Kennedy et al. 2019) along with less penetrant phenotypes such as seizures, microcephaly and autism spectrum disorder. Most of the pathogenic mutations characterized to date are protein-truncating mutations that occur throughout the length of the gene, but are most commonly observed in the last exon that makes up over half of the gene. Missense mutations to the enzymatic histone acetyltransferase domain or highly conserved C-terminus regions have also been associated with ARTHS, however, the phenotype is less-severe than the protein-truncating mutations (Kennedy et al. 2019). The majority of patients with late-truncating mutations in KAT6A, located beyond exon 16, have more severe phenotypes compared to patients with early-truncating mutations in exons one through fifteen (Kennedy et al. 2019).

*KAT6A* is part of the orthologous MYST family of highly conserved histone acetyltransferase (HAT) genes (Thomas and Voss 2007) that modulates gene expression through deposition of acetyl groups on the histone tail, which increases chromatin accessibility at specific loci and allows for spatial arrangement of transcription factors and transcriptional machinery local control of gene expression (Nava and Arboleda 2023). Recent studies in mammalian model systems have demonstrated that KAT6A acetylates specific lysine residues on histone H3, including lysine 9 (Voss et al. 2009; Arboleda et al. 2015), and lysine 23 (Yan et al. 2017, 2020). Lower abundance modifications, such as propionyl modifications, have also been described (Yan et al. 2020) but are near impossible to validate due to limited antibody specificity and abundance. One of the key questions that remains is the extent to which *KAT6A* coordinates and directs transcriptional activation of specific genes and pathways in human cells, especially in the context of ARTHS.

The *KAT6A* gene exhibits significant structural conservation across eukaryotes indicating a shared biological functionality throughout evolution. In model systems, the KAT6A homologs have been shown to influence gene silencing (Reifsnyder et al. 1996), oocyte polarity (Huang et al. 2014), hematopoiesis (Katsumoto et al. 2006; Genais et al. 2020), and neuroblast proliferation (Reifsnyder et al. 1996; Scott et al. 2001). Tissue-specific *Kat6a^-/-^* have demonstrated a key role of KAT6A in adult hematopoietic stem cell maintenance (Perez-Campo et al. 2009; Sheikh et al. 2016) and in craniofacial patterning through nitric oxide signaling and histone acetylation (Kong et al. 2014; Vanyai et al. 2019). In mice, knockout of *Kat6a* does not globally decrease H3K9 acetylation but leads to hypo-acetylation at specific loci, and reduced expression of *Hoxa* and *Hoxb* gene clusters compared to controls driven by hypoacetylation of the transcriptional start sites controlling expression of *Hoxa4*, *Hoxb3* and *Hoxb4* (Hibiya et al. 2009; Voss et al. 2009). The *Hox* gene clusters play a vital role on body patterning during early developmental timepoint (Mallo et al. 2010; Hubert and Wellik 2023) and in development of the central nervous system (Nolte and Krumlauf 2013; Philippidou and Dasen 2013).

In this study, we perform an epigenomic and transcriptomic assessment on fibroblasts from 8 ARTHS patients and 14 control individuals. All patients harbor a genetic diagnosis of ARTHS and have a pathogenic *KAT6A* mutation that results in premature truncation of the protein (**Table S1**). Our analysis identified 60 genes that were differentially expressed in ARTHS-patient dermal fibroblast samples as compared to controls. Our multi-omic dataset demonstrates that protein truncating mutations in *KAT6A* cause differential chromatin accessibility, histone acetylation and transcriptomic changes control posterior *HOXC* genes (*HOXC10*, *HOXC11*, *HOTAIR*, *HOXC-AS3* and *HOXC-AS2*). Furthermore, a mass spectrometry based assay on histone extracts identified novel histone PTMs on the Histone 2A and Histone 3 that are disrupted by *KAT6A* mutations. Our study provides valuable insights into the gene regulatory mechanisms that are perturbed in primary samples of Arboleda-Tham syndrome.

## RESULTS

To study the direct effect of *KAT6A* mutations on gene expression in ARTHS patients, we established fibroblast cell lines from ARTHS patients (n=8) and unaffected individuals (n=14). The pathogenic mutations in the patients disrupted the KAT6A protein between amino acid 379 and 1551 with seven of the eight mutations being *de novo truncating* mutations and one being a *de novo* missense change in a highly-conserved region of *KAT6A* (**Figure 1A, Figure S1)**. The clinical phenotypes of these eight patients were previously reported (Arboleda et al. 2015; Tham et al. 2015; Kennedy et al. 2019).

**Figure 1.**
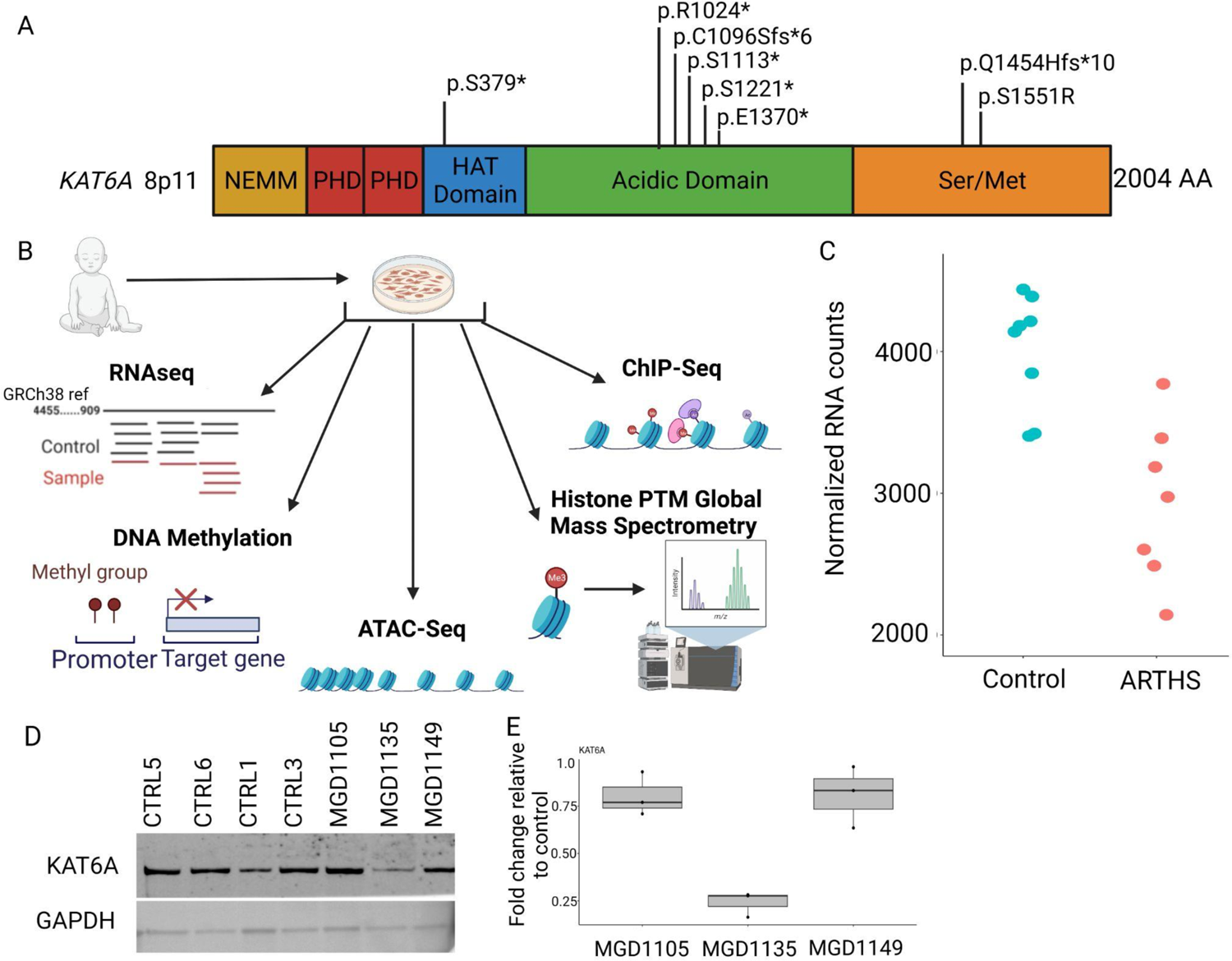
*KAT6A* mutations in patient-derived dermal fibroblasts undergo multi-omic profiling. A) KAT6A protein structure highlighting important domains and location of the eight pathogenic mutations included in our current study. B) In this study, ARTHS fibroblasts were subject to transcriptomics (RNA-seq), Histone post-translational modification profiling by mass spectrometry, DNA Methylation, ATAC-seq and ChIP-Seq for histone PTMs. C) Assessment of *KAT6A* mRNA expression from transcriptome profiling show slightly decreased levels of *KAT6A* expression in ARTHS patients compared to controls. D) Western blot of KAT6A protein show decreased protein only in MGD1135 which harbors an early-truncating nonsense mutation at p.S379*. E) Quantification of western blots show an 80% decrease in protein levels in MGD1135.

To understand how these *KAT6A* mutations disrupt the epigenome and RNA expression, we performed epigenomic and transcriptomic assays (**Figure 1B**). Previous studies have pointed towards disruption of histone post-translational modification of specific genomic loci rather than a genome-wide decrease of specific histone marks, we performed genome-wide epigenetic assays to assess chromatin accessibility (ATAC-seq (Corces et al. 2017), chromatin Immunoprecipitation of H3K9 and H3K23 acetylation, DNA methylation using Epic 850K array and, RNA sequencing. We also performed a histone-targeted mass spectrometry assay to identify novel histone PTMs that might be regulated by KAT6A (**Figure 1B, Table S1**). Representative samples were assessed for each assay.

We first assessed the RNA and protein levels of KAT6A in our dermal fibroblast lines. We found a modest decrease in KAT6A RNA (**Figure 1C**) which was not statistically significant. At the protein level, we did not see an appreciable difference in two of the three mutations. We did notice that our early truncating mutation patient, MDG 1135, that has a nonsense mutation at amino acid position 379, showed a 80% decrease in protein levels and is the earliest truncating mutation line we tested (**Figure 1D, 1E, Figure S2**).

### *KAT6A* mutations lead to modest changes in the transcriptional landscape

We next performed RNAseq analysis from eight controls and seven patients and assessed differential gene expression between the cases and controls. As expected, the ARTHS cases and controls separated out in unsupervised clustering (**Figure 2A**) across the 60 genes that were identified as significantly differentially expressed (p_adj_ < 0.05) between control and ARTHS fibroblasts. The significantly differentially expressed genes were predominantly protein coding genes (46/60) but a smaller subset were long noncoding RNA (lncRNA) (9/60) (**Figure 2B, Table S2**). An equal number of genes were upregulated and downregulated in patients with ARTHS (50%, 30/60).

**Figure 2.**
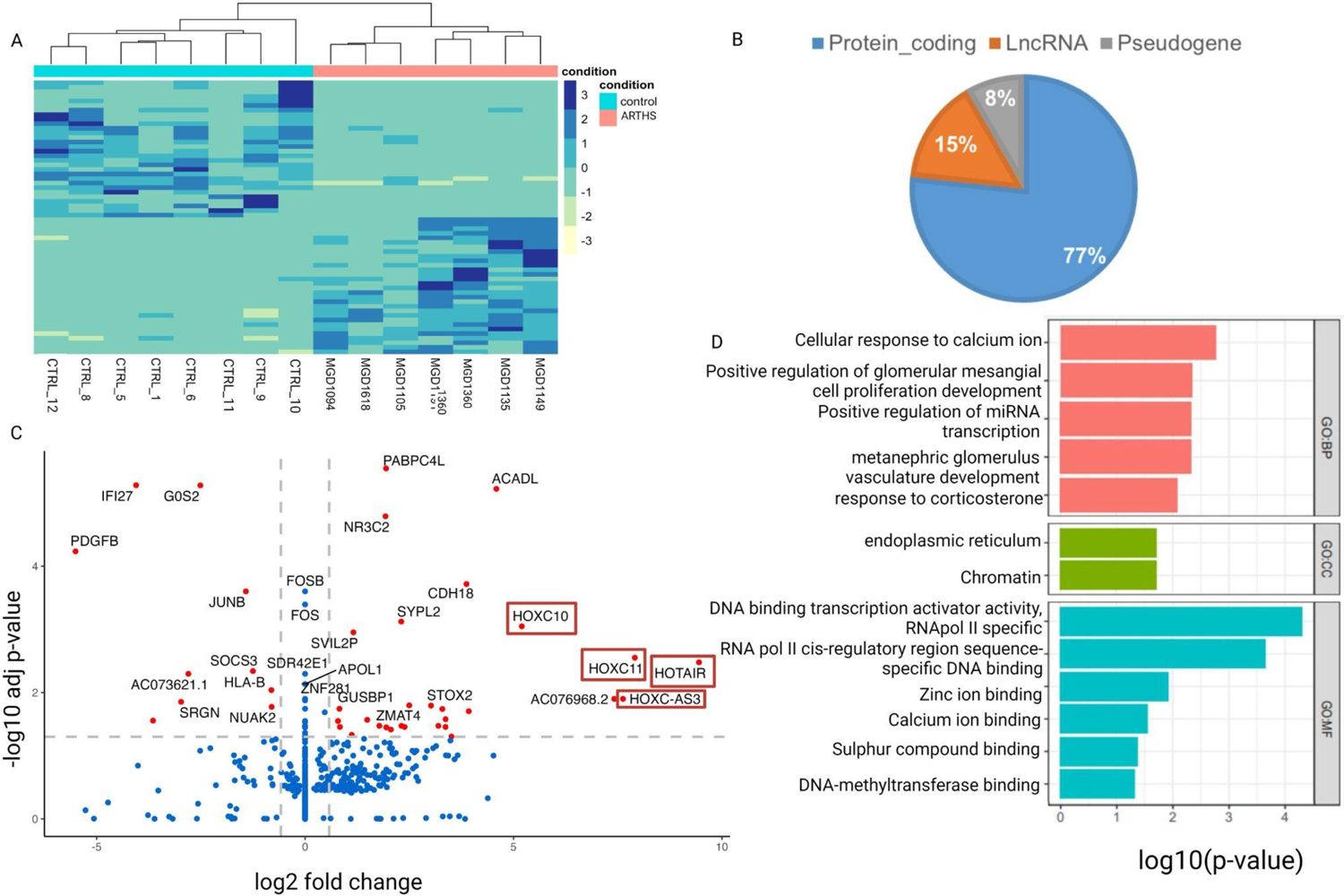
*KAT6A* mutations cause upregulation of the posterior HOXC cluster in ARTHS fibroblasts. A) Heatmap of 60 significantly differentially expressed genes (DEGs) in ARTHS fibroblast compared with controls. B) The gene type distribution of the DEGs show most of the RNAs identified represent protein-coding genes (blue). C) Volcano plot of significant DEGs in ARTHS fibroblast compared with control samples highlight 40 genes that have an absolute fold-change of 1.5 and p-adjusted of less than 0.05 (red). All other genes are represented in blue. D) Gene Ontology bar graph for the differentially expressed genes for biological processes (BP), cellular components (CC) and molecular function (MF).

We noted that 4 of the 5 genes with the highest fold change in ARTHS fibroblasts were a set of developmentally regulated genes that belong to the HOX gene cluster. Specifically, this subset of genes include: *HOXC11* (p_adj_=2.8e-3, log2FC=7.9), *HOXC10* (p_adj_=8.9e-4, log2FC=5.2), *HOXC-AS3* (p_adj_=2.0X1-3, log2FC=7.2), *HOTAIR* (p_adj_=3.0e-3,log2FC=9.4) and *HOXC-AS2* (p_adj_=1.8e-2, log2FC=3.3) (**Figure 2C, Table S2**). The HOXC gene cluster is located on chromosome 12 and, like all the other three *HOX* gene clusters, oriented in a specific order that corresponds to their spatial-temporal expression during early embryonic patterning (Wellik 2007). The redundancy among some of the genes within each HOX gene cluster suggests that they have overlapping functions and that combinatorial expression drives the early patterning events (Pearson et al. 2005).

Although fibroblasts are not necessarily representative of the most affected cell-type in ARTHS, we are able to identify differentially expressed genes highlighting a potential role for KAT6A in differentiated cells. In the ARTHS fibroblasts RNAseq, we identified transcriptionally dysregulated genes with established neural-related function. Here we highlight 5 neural-related genes (i.e. *CDH18*, *PITX1*, *PCP4*, *PABPC4L*, *PKNOX2*) that we discovered are significantly upregulated in ARTHS fibroblast compared to unaffected controls. *Cadherin 18* (CDH18) (p_adj_=4.8e-8, log2FC=3.8) is expressed in neurons and implicated in neuropsychiatric disorders (Redies et al. 2012). *Paired like homeodomain 1* (*PITX1*) (p_adj_=1.0e^-^4, log2FC=3.5) deletions in humans lead to lower limb deformities and other skeletal anomalies (Klopocki et al. 2012) but the effect of overexpression of *PITX1* remains unknown. *Purkinje cell protein 4* (*PCP4*) (p_adj_=3.3e-5, log2FC=3.4) is another upregulated gene that is known to modulate calmodulin binding activity in neuronal cells (Mouton-Liger et al. 2011). Poly(A) binding protein, cytoplasmic 4-like (PABPC4L) (p_adj_=8.9e-4, log2FC=5.2) is an RNA binding protein that has recently been shown to be involved in familial atypical Parkinsonism (Aslam et al. 2019). PBX/knotted 1 homeobox 2 (*PKNOX2*) (p_adj_=3.67e-8, log2FC=3.6) is also upregulated. PKNOX2 targets the PBX proteins, and is hence thought to regulate tissue specific protein expression (Imoto et al. 2001). PKNOX2 has also been identified as a candidate gene for schizophrenia in two large studies (Wang et al. 2012).

Of the genes that are transcriptionally downregulated, we show that *KAT6A* mutations are associated with significant downregulation of *platelet derived growth factor subunit B* (*PDGFB*) (p_adj_=1.25e-8, log2FC=-5.5) in ARTHS fibroblasts. Endothelial cell-specific deletion of *Pdgfb* leads to vascular defects across highly vascular organ systems such heart and kidney (Levéen et al. 1994; Bjarnegård et al. 2004). This is consistent with earlier reports of prenatal lethality due to failure of blood development in *Kat6a^-/-^* knockout mice.

Gene Ontology enrichment of the differentially expressed genes, identified critical pathways and cellular components. The differentially expressed genes were enriched for roles in chromatin, which confirmed the quality of our data given the known role of *KAT6A* as a histone acetyltransferase. We also identified significant enrichment for DNA-binding activity, and calcium and zinc ion binding (**Figure 2D**) that are related to transcriptional regulation. The enriched GO terms were driven by the inclusions of *JUNB*, *FOS*, *FOSB*, *HOXC10*, *HOXC11* and *EGR1* genes (**Table S3**).

### *KAT6A* mutations alter chromatin accessibility and are associated with differential gene expression

Since KAT6A protein belongs to the class of chromatin modifiers, we next asked whether the differences in gene expression might be associated with changes in chromatin accessibility. We performed ATACseq on ARTHS and control fibroblasts to see if *KAT6A* mutations are associated with local disruption of chromatin state. Our analysis identified a total of 514 significantly differentially accessible regions, of which 287 peaks (55.8%, 287/514) correspond to more open regions in ARTHS fibroblasts and 227 peaks (44.2%, 227/514) that were more closed in ARTHS fibroblasts (**Figure 3A**). Of all the peaks identified, 56.9% (292/514) of the peaks correspond to intergenic regions, 189 peaks correspond to gene body regions (5’UTR = 5/514, 3’UTR = 3/514, Intron = 181/514), and 3.8% (20/514) peaks correspond to promoters of known genes. These 514 differentially accessible peaks correspond to 414 unique genes (**Figure 3B**). In this dataset, of all the closest genes identified, 59% (242/414) are protein-coding, and 38% (156/414) are ncRNAs (non-coding RNAs) (**Table S4**).

**Figure 3:**
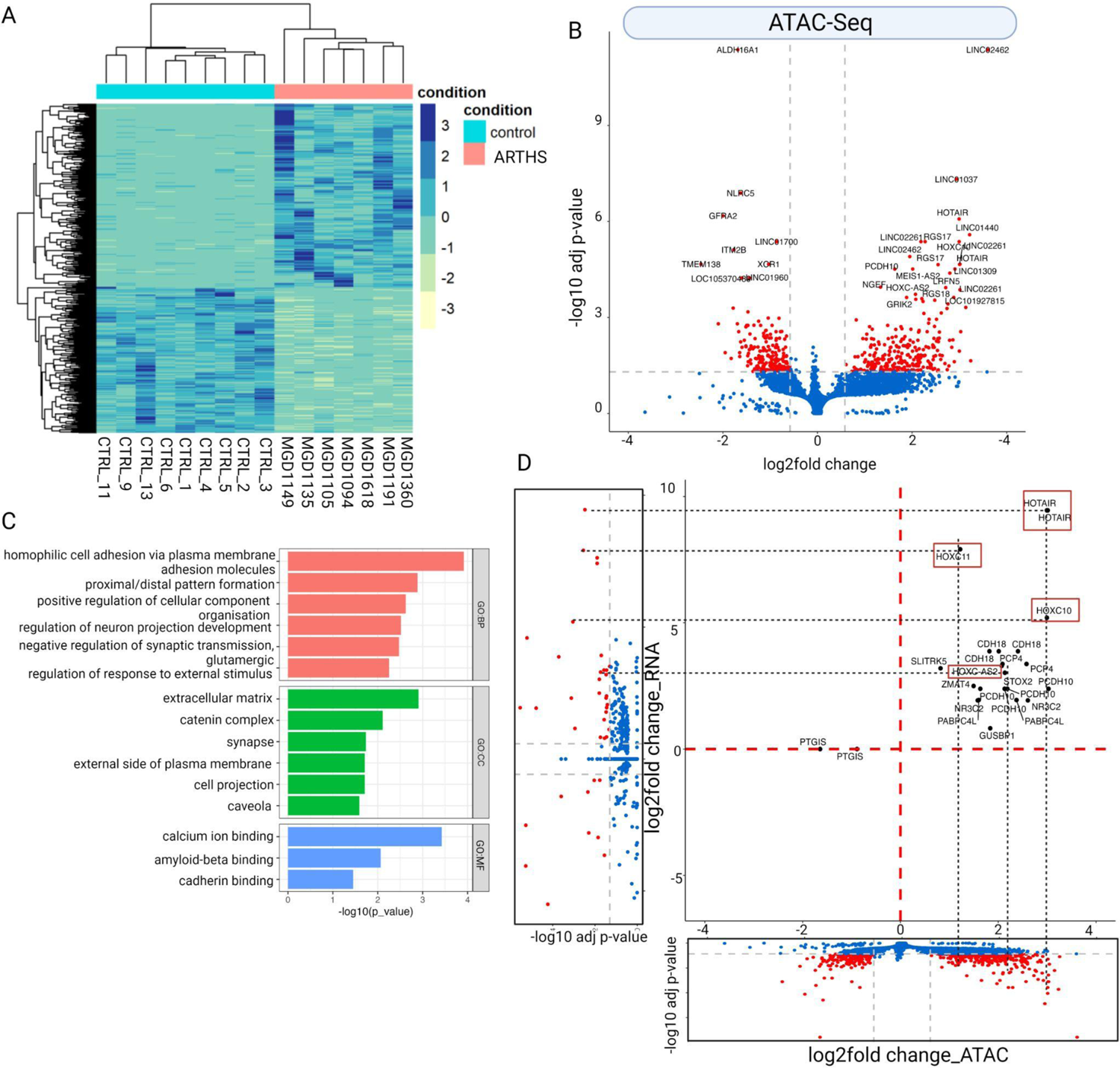
*KAT6A* mutations show that differential chromatin accessibility drives 23% of genes that are differentially expressed: A) Heatmap depicting 514 differentially accessible ATAC peaks between patient and control samples B) Volcano plot highlighting 395 genes that have an absolute fold-change of 1.5 and p-adjusted of less than 0.05 (red). All other genes are marked in blue. C) Significant GO term distribution of the SigDEGS highlighting biological processes of neuron projection and synaptic transmission. D) Correlation between ATAC and RNAseq data. It highlights the expression change and chromatin accessibility being changed on *HOXC* genes.

We next wanted to investigate whether differentially accessible regions and the genes associated with these open chromatin regions had common features or harbored genes that drove specific cellular signaling pathways. We used GeneOntology (GO) (Raudvere et al. 2019) enrichment analysis on the list of 414 genes located closest to differentially accessible regions, and we identified enrichment for cellular component organization, neuron projection, synaptic regulation, RNA and protein binding (**Figure 3C, Table S5**).

Finally, we asked whether there was a correlation between changes in the chromatin accessible regions with gene expression changes. To address this, we asked whether genes that display increased chromatin accessibility also have higher RNA expression, and vice versa, in ARTHS fibroblasts. We performed correlation between all the genes that were differentially accessible and differentially expressed in ARTHS as determined by the ATACseq and RNAseq analysis, respectively. We showed that several genes in the *HOXC* cluster are differentially expressed (**Figure 2C**) and we also find that these same genes have differential chromatin accessibility in our ATACseq data (**Figure 3D**). There is a broad accessibility to the chromatin around the HOXC gene cluster and is correlated with increased gene expression in ARTHS samples (*HOTAIR* (p_adj_= 2.16e-5, log2FC=3.01), *HOXC10* (p_adj_=4.27e-6, log2FC=2.9), HOXC-AS2 (p_adj_=1.8e-4, log2FC=2.07), *HOXC9* (p_adj_=2.7e-4, log2FC=2.07), HOXC11 (p_adj_=1.2e-2, log2FC=1.22) (**Table S4**). Overall we find that *KAT6A* mutations lead to localized specific changes to chromatin accessibility.

### *KAT6A mutation*s lead to an differential H3K9 and H3K23 acetylation at specific genomic loci

Previous studies have identified several histone 3 lysine acetylations that were disrupted in the presence of *KAT6A* mutations, including H3K9 (Voss et al. 2009; Arboleda et al. 2015) and H3K23 (Huang et al. 2014; Lv et al. 2017; Yan et al. 2020). We first confirmed that global histone levels were not changed by *KAT6A* mutations by performing western blots using antibodies specific to H3K9ac and H323ac and did not identify any significant global differences between ARTHS and control samples (**Figure 4A, 4B and 4C, Figure S3A and S3B**).

**Figure 4.**
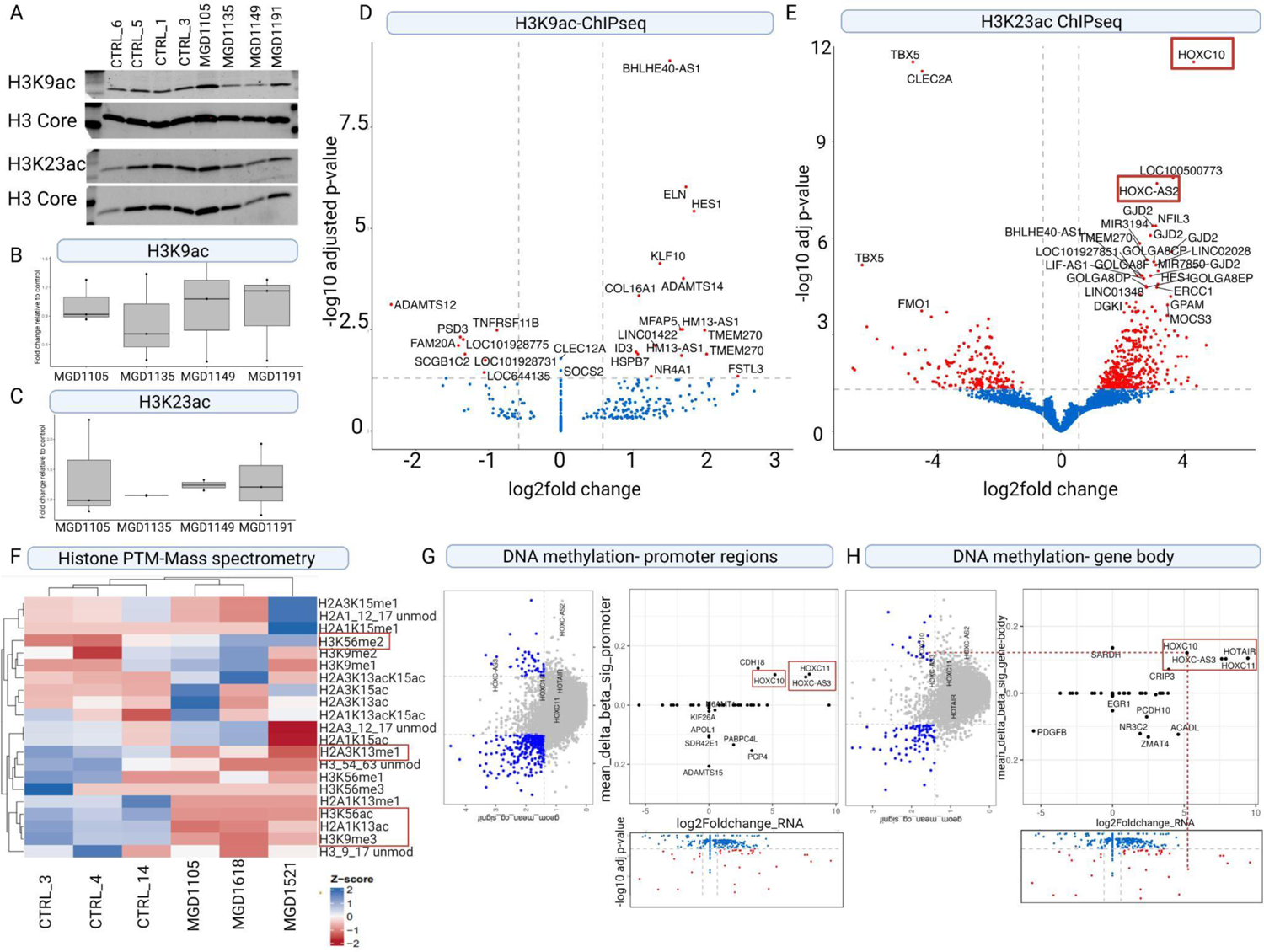
Histone post-translational modifications and DNA-methylation are disrupted in ARTHS fibroblasts around posterior *HOXC* genes. Western blot of global histone acetylation for A) H3K9ac and H3K23ac are performed compared to H3 core B) Relative quantification of the westerns does not show significant diffrences in global PTM levels of B) H3K9ac; and C) H3K23ac. Volcano plot showing the genes corresponding to significantly differentially accessible peaks associated with D) H3K9 acetylation and E) H3K23 acetylation. Genes showing significant differences (padj <0.05 and abs(log2FC)>0.58 are highlighted in red. All other genes are highlighted in blue. F) Heatmap of the normalized abundance of single histone PTMs for peptides that were detected and identified as having at least one histone PTM that is significantly different between ARTHS fibroblast and control samples as determined by mass-spectrometry. Histone PTMs that were found to be significantly different (p<0.05) between are boxed in red. Correlation between DNA methylation on G) promoter regions and RNAseq data and H) Gene-body regions and RNAseq data. We observe differential methylation across CpG islands being changed on *HOXC* genes.

Given this, we asked whether there were specific genomic loci where H3K9 and H3K23 were disrupted. We performed Chromatin Immunoprecipitation sequencing (ChIPseq) to identify genomic loci that have disrupted H3K9Ac (**Figure 4D, Figure S3C, TableS6**) and H3K23Ac (**Figure 4E, Figure S3D, Table S8**) binding in ARTHS samples. The direct number of peaks identified in H3K9Ac was limited to 27 peaks in 24 unique genes. Of these 41% (10/24) were hypoacetylated and 58% (14/24) were hyperacetylated. However, H3K23ac ChIPseq identified 603 significantly differential accessible peaks which correspond to differentially acetylated regions at the H3K23 residue (**Figure 4E, Figure S3D, Table S8**). These 632 peaks correspond to 403 unique genes. A gene ontology enrichment for the genes closest to the peak (**Figure S4E and S4F, Table S7 and S9**) again showed enrichment for terms like RNA and protein binding in the H3K23ac dataset. Two loci that were significantly bound to H3K23 acetylation in ARTHS were directly located in the posterior *HOXC* gene cluster that were differentially expressed and showed differential chromatin accessibility. Together our data suggests that H3K23 acetylation drives the increased chromatin accessibility around the HOXC cluster and increased gene expression that is observed in ARTHS fibroblasts.

### Histone mass spectrometry identifies differential histone post-translational modifications in ARTHS samples

Given that *KAT6A* functions as a histone acetyltransferase, we wanted to investigate the repertoire of histone residues that might be acetylated by the *KAT6A* mutation and those that might be indirectly affected by changes in KAT6A’s acetylation pattern. Previous studies have focused on established marks for which there are well-curated antibodies that can quantitatively measure acetylation levels. However, there are many more acetylation marks across H2, H3 and H4 that have not been comprehensively queried in these patients. Furthermore, assessment of not just acetylation marks but also changes to alternative post-translational modifiers that might represent indirect events shed light on the combinatorial role of different histone post translational modifications.

To address this question we performed mass spectrometry (MS) on histone extracts from 3 ARTHS and 3 control fibroblast samples to identify differential histone modifications in ARTHS relative to unaffected controls (**Table S10**). The MS data did not show any global changes to the H3K9ac (p=0.288, log2FC = 0.362) or H3K23ac (p=0.356, log2FC= 0.253) marks in ARTHS.

However, as shown in **Figure 4F**, we did observed a significant depletion of several novel histone marks that have not previously been associated with KAT6A: H3K56ac (p=0.0086, log2FC= − 4.95), H3K9me3 (p=0.0046, log2FC= −0.196), H2A.1K13ac (p= 0.0043, log2FC= −0.649), and H2A.3K13me1 (p= 0.0212, log2FC= −0.261). We also discovered significantly increased levels of H3K56me2 (p=0.0273, log2FC= 1.23) (**Figure 4F)**. In yeast, H3K56ac impacts global transcriptional activation (Rufiange et al. 2007; Topal et al. 2019), while in mammals H3K56ac plays a role in cell division and chromosomal organization by depositing this mark in newly formed chromatin (Fang et al. 2022). Histone-mass spectrometry is a powerful tool for comprehensively identifying novel PTMs disrupted by genetic mutations in chromatin modifiers.

### ARTHS samples have hypomethylated CpGs as compared to controls

Finally, we obtained previously published DNA-methylation data from ARTHS fibroblasts and controls (Bondhus et al. 2022). Our data indicates that *KAT6A* mutations lead to altered epigenome to regulate the expression of specific gene sets. To gain a more in-depth understanding of this regulation, we used the DNA methylation array on bisulfite converted DNA from five ARTHS and four control cell lines to identify differentially methylated CpGs and regions. DNA methylation array data shows that the CpG sites are more hypomethylated in ARTHS samples as compared to controls (**Figure 4G** and **4H**, **Figure S4A,S4B** and **S4C**). With this dataset we observed a modest hypermethylation in the gene body region of the *HOXC* genes was correlated with higher transcription (Figure 4H). The role of hypermethylated CpGs in the promoter regions have been well established and are associated with gene silencing, however, there have been few studies that have shown that the methylated CpGs in the gene body have an opposite effect and are associated with active transcription (Hellman and Chess 2007; Ball et al. 2009; Laurent et al. 2010).

### Multi-omic data integration shows that *KAT6A* mutations cause overexpression of the posterior *HOXC* cluster genes

Our multi-omics data analysis identified differential chromatin accessibility and acetylation around posterior *HOXC* cluster genes (**Figure S5A and S5B**). Since *HOX* genes play spatiotemporal roles during development and maintenance of specific cell type, and they manifest a collinear expression pattern, we wanted to see how the chromatin accessibilty, epigenome and RNA expression is maintained around the entire HOXC clusters. We created coverage plots mapped around the entire *HOXC* gene locus, we observed no changes to the anteriorly parts of the HOXC cluster, across ATACSeq, ChIPSeq for H3K23ac and gene expression (**Figure 5A**). We see increased accessibility, acetylation and expression of the posterior *HOXC* genes (*HOXC11*, *HOXC10*, *HOTAIR*, *HOXC-AS2* and *HOXC-AS3*) (**Figure 5B**) and DNA hypermethylation at CpG sites in the posterior HOX genes (**Figure 5C)**. Our study indicates that the *KAT6A* mutations have an important impact on transcriptional regulation of the posterior *HOXC* genes.

**Figure 5.**
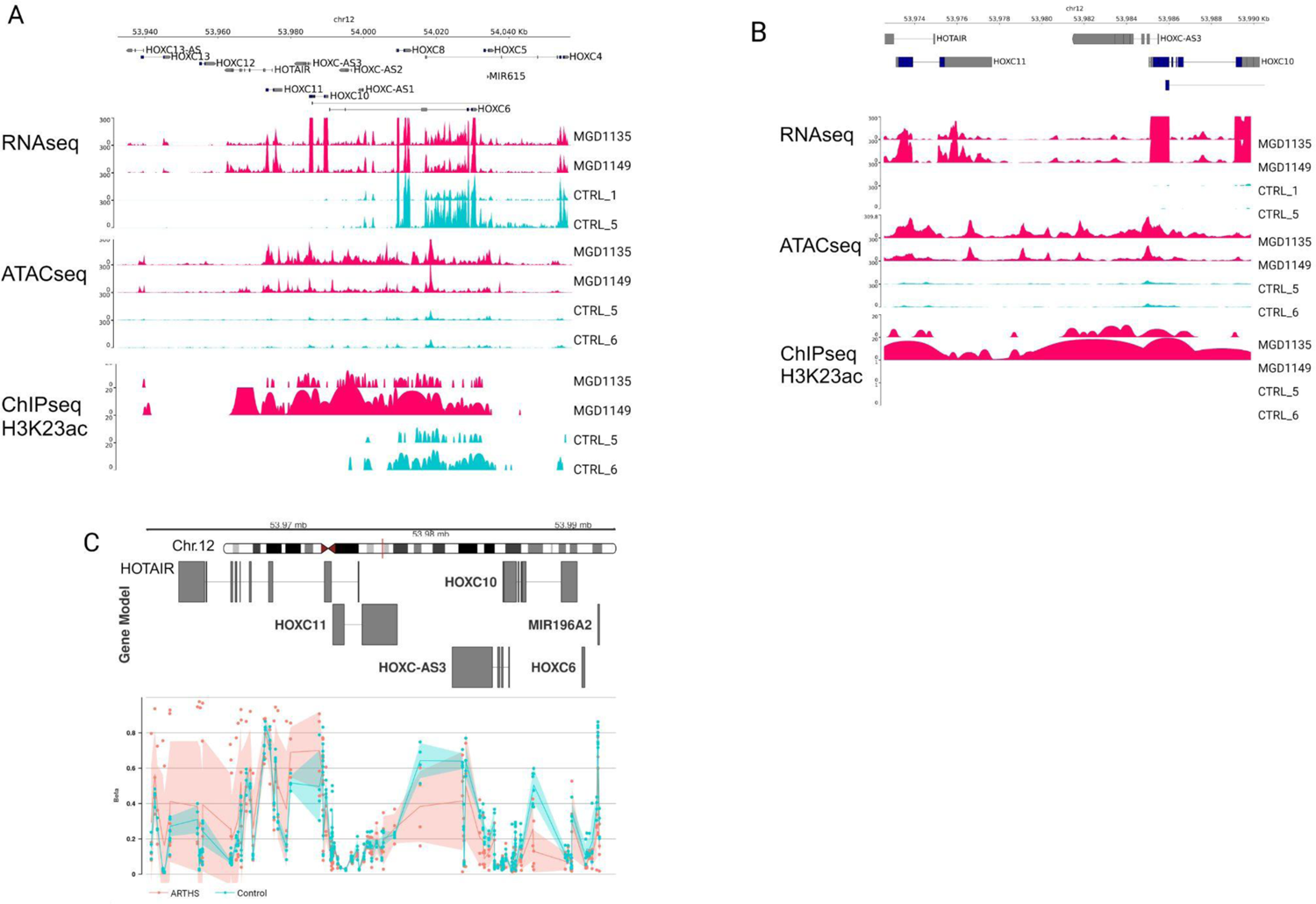
ARTHS mutations regulate the expression of posterior *HOXC* genes through H3K23 acetylation. Multi-omic epigenomic and transcriptomics data from ARTHS samples compared to controls shows clear regulation of posterior *HOXC* cluster genes. A) Coverage tracks for RNA-seq, ATAC-seq and ChIP-seq across the entire *HOXC* cluster. B) Zoomed-in coverage of RNA-seq, ATAC-seq, ChIP-seq across the posterior *HOXC* genes (*HOXC10*, *HOXC11*, *HOTAIR*, *HOXC*-*AS2*, *HOXC-AS3*) C) Differential CpG methylation pattern across the posterior *HOXC* cluster.

The knowledge gained from this multi-omics data integration coupled with the current literature around *KAT6A* and *HOX* genes indicate that different types of *KAT6A* genetic mutations in model organisms have differential effects on HOX gene expression. Overexpression of HOX genes have been associated with cancer(Kim et al. 2019; Fang et al. 2021) and motor neuron defects(Misra et al. 2009). We show that in our model, which is unique in that it leverages human fibroblasts from ARTHS patients, truncating *KAT6A* mutations epigenetically upregulates *HOXC* gene cluster expression. Our study suggests that *KAT6A* mutations in ARTHS fibroblasts cause HOX overexpression which likely contributes to gross motor phenotypes in ARTHS individuals, via mechanisms similar to transgenic overexpression of Hox gene in model organism studies

## DISCUSSION

Our multi-omic study of primary ARTHS fibroblast samples quantitatively assayed the epigenomic-driven changes in gene expression across different pathogenic KAT6A mutations. For the first time, we precisely identify the genomic loci and epigenetic marks that are transcriptionally dysregulated in primary ARTHS samples as a result of pathogenic KAT6A mutation through the systematic integration of our RNAseq, ATAC-seq, ChIP-seq, and Histone-PTM-specific mass spec data. Most notably, we identified that differential chromatin accessibility and H3K23 acetylation within the *HOXC* gene cluster drove extremely high expression of HOXC10, HOXC11, HOTAIR, HOX-AS3 and HOXCAS2. Moreover, our unbiased Histone PTM proteomics approach allowed us to identify depletion of acetylated histone marks at H3K56 and H2A.1K13 in ARTHS fibroblasts compared to controls, which suggests KAT6A may regulate the acetylation status of these previously undescribed histone marks in the context of ARTHS. Taken together, our multi-omic profiling of primary ARTHS fibroblasts allowed us to glean novel insights into the gene regulatory biology controlled by KAT6A.

The *KAT6A* gene is part of a broader group of genes that, when mutated, cause chromatinopathy syndromes due to disruption of epigenome regulation (Nava and Arboleda 2023). Chromatinopathies affect multiple tissues and organ systems, making the study of patient primary cells (e.g. fibroblast and blood) an invaluable resource for scientific discovery (Nava and Arboleda 2023). Since *KAT6A* functions as an epigene, we can measure the downstream molecular effects through multi-omic measurements of downstream markers such as: histone PTMs, localization of histone PTMS, chromatin accessibility, and gene expression. To date, this represents the first multi-omic study of ARTHS in which we leverage patient-derived cells to directly interrogate the role of KAT6A on the epigenome and gene expression.

### *KAT6A* mutations disrupt *HOXC* gene cluster that is associated with lumbar motor neuron development

The *HOX* genes are an evolutionarily conserved class of genes, initially described in drosophila, that play a significant role in differentiation and anterior-posterior body patterning (McGinnis and Krumlauf 1992)(Khan et al.,Perinatal and Developmental Epigenetics Volume 32 in Translational Epigenetics 2023, Pages 71-113). In vertebrates, the *hox* gene cluster underwent gene-level duplications resulting in four clusters of *HOX* genes located on different chromosomes (Sánchez-Herrero 2013; Nolte et al. 2015). The *HOX* cluster genes are expressed in a co-linear fashion and coordinate temporal and spatial patterns during development. The clusters allow for some redundancy within essential processes and fine tuning of the process by which the three germ layers differentiate into cell, tissues and organ systems in three dimensional space (Kmita and Duboule 2003; Dasen and Jessell 2009).

We and others have identified disrupted expression of HOX genes in various model systems and tissues where the *KAT6A* gene is mutated. In normal development, mouse studies have shown that at E9.5, *Hoxa* and *Hoxb* are localized to the hindbrain while *Hoxc* and *Hoxd* are localized to the spinal column at E12.5. *Hoxc10* and *Hoxc11* are restricted to the lumbar region of the spinal column and are essential to hind-limb motor neuron development (Hostikka and Capecchi 1998; Wellik and Capecchi 2003; Wu et al. 2008; Dasen and Jessell 2009). The upregulation of posterior *HOXC* cluster genes in our data may have an impact on the early development of motor neurons, potentially contributing to the hypotonia and gross motor delay observed in ARTHS patients. In our dataset, we identified the upregulation of five genes from the HOXC cluster: *HOXC10*, HOXC11, *HOXC-AS3*, *HOXC-AS2*, and *HOTAIR*. Interestingly, previous studies have primarily associated defects in motor neuron development with the absence of Kat6a/Moz/Myst3 and, consequently, the Hox cluster gene.

While much attention has been focused on *HOX* gene deletions, the effect of *HOX* overexpression in normal development remains unclear. Notably, a study in a chicken embryo model demonstrated that overexpression of *Hoxd10* leads to a reduction in the total motor neuron population and shifted the population towards increased lumbar motor neurons (Misra et al. 2009). Similarly, in mice, the duplication of *Hoxd11* results in altered development of axial skeletal tissue (Boulet and Capecchi 2002). Overexpression of the *HoxC* cluster genes has not been independently studied but would be required to link developmental HoxC overexpression to a specific aspect of the ARTHS phenotype in patients.

While our findings in ARTHS patients did not reveal changes to KAT6A expression at the transcript or protein level, we believe that presence of the truncated protein product causes mistargeting of the KAT6 complex and disruption of *HOXC10 and HOXC11* gene expression, in ARTHS fibroblasts. Subtle shifts in cellular populations can disrupt normal neural connectivity leading to some of the clinical features that are characteristic of ARTHS. Further studies in complex systems that directly link the KAT6A mutations to these loci and histone marks would be required to prove the direct effects of KAT6A mutations on these critical regions.

### *KAT6A’s* effect on HOXC10 gene expression is conserved across different cell types derived from ARTHS patients

The diversity enabled by epigenome organization is a major mechanism that allows a single genome to give rise to thousands of diverse cell types in the human body (Nava and Arboleda 2023). While the genome remains the same, the epigenome coordinates the structure and conformation of specific genomic loci to direct differentiation of each cell type. *KAT6A* is an epigene, which is defined as a protein-coding gene that regulates or affects the epigenome structure and function (Nava and Arboleda 2023). Therefore, disruption of *KAT6A* function is known to affect acetylation of specific histone marks in the KAT6 complex (Zu et al. 2022). But how the KAT6 complex is directed towards specific genomic loci for acetylation during growth and differentiation is not completely understood, especially in the context of human biology (Nava and Arboleda 2023). Strikingly, upon comparing the genes that were differentially expressed in our ARTHS fibroblast data to genes that were differentially expressed in ARTHS cerebral organoids--we uncovered that *HOXC10* was significantly upregulated in ARTHS across both independent RNAseq datasets in comparison to unaffected individuals (Nava et al. 2023). This suggests that some of the *HOXC* cluster genomic targets of KAT6A are conserved across different cell types in ARTHS. Moreover, upon comparison to ARTHS cerebral organoid study, we found there were other *HOX* genes from other clusters (e.g. *HOXA*, *HOXB*, *HOXD* clusters) that were significantly overexpressed in ARTHS cerebral organoids but not in our ARTHS fibroblast data (Nava et al. 2023). This suggests that the HOX cluster genomic targets of KAT6A vary across different human cell types, which may reflect differences in post-translational or post-transcriptional regulation of HOX cluster genes in ARTHS. Studies in mice, zebrafish and medaka fish have shown that *KAT6A* deletion decrease expression of specific *HOX* gene cluster expression(Miller et al. 2004; Hibiya et al. 2009; Voss et al. 2009) indicating a potentially dual and cell-type and organism specific effect of KAT6A regulation of *HOX* gene clusters. Our study is the first that demonstrates genetically driven upregulation of posterior *HOXC* genes.

### Global effects on specific histone acetylation modifications due to *KAT6A* mutations

KAT6A is a lysine acetyltransferase enzyme that is involved in lysine acetylation of Histone 3 N-terminal tail (Dreveny et al. 2014; Lv et al. 2017). However, most studies are done on a subset of possible acetylation markers, based on the availability of histone-mark specific antibodies. In ARTHS fibroblasts, global histone acetylation of H3K9 or H3K23 was not decreased and ChIPseq studies identified a moderate number of loci that showed differential H3K9 or H3K23 acetylation (**Figure 4D and 4E**) several of which correlated with the differential gene expression. We performed the first unbiased histone mass spectrometry assays to comprehensively explore all histone PTMs that are directly or indirectly altered by *KAT6A* mutations. We believe mass spectrometry assay has broad potential to explore global changes across histone modifications. This, coupled with ChIP-seq to assess localization of these histone marks, our studies show that the commonly assayed H3K9ac or H3K23ac marks did not show significant changes to global acetylation by mass spectrometry or significant changes on the localization of these marks (**Figure 4A-E**). Our study has identified two novel marks that show global depletion upon KAT6A mutations in fibroblasts: H3K56ac and H2AK13ac.

The role of H3K56ac has been well established in yeast, where it plays crucial roles in chromatin remodeling and DNA repair (Shuttleworth 1991; Xu et al. 2005). Recent studies in mammals also support the role of H3K56ac in chromatin stability and DNA damage and repair (Yuan et al. 2009). These are consistent with earlier studies demonstrating KAT6A’s role in DNA damage and repair (Rokudai et al. 2013). H3K56ac has been shown to facilitate the binding of certain pluripotency markers or transcription factors in mouse embryonic stem cells (Tan et al. 2013). Our mass spectrometry data analysis also shows changes in H2A13Kac in ARTHS samples as compared to control. H2AK13ac has not been extensively studied and more in depth investigations on the gene regulatory effect of H2AK13ac are required, however it is believed that the H2A and H2B play essential roles in chromatin organization.

Our findings expand on previous studies showing that *KAT6A* mutations disrupt *HOX* gene expression but that the gene regulation by KAT6A is likely to be cell-type and tissue-specific. Furthermore, our newly described connection to H3K56 acetylation and H2AK13 acetylation highlights the critical role of unbiased approaches in identifying associated histone marks that are globally disrupted by epigenes. To gain a comprehensive understanding of the role of *KAT6A* mutations in human development and disease, further in-depth studies using relevant human tissue types or model systems are warranted. Such investigations will contribute to unraveling the precise mechanisms underlying how *KAT6A* mutations disrupt normal development and disease.

## METHODS

### Cell Culture

Skin punch biopsies were performed on patients with a confirmed pathogenic KAT6A mutation and on control individuals under IRB#11-001087 approved by the UCLA Institutional Review Board. Skin punch biopsies were performed and processed to create dermal fibroblast lines in the UCLA Pathology Research Portal. Control neonatal, GM01651 and GM00323 fibroblast lines were obtained from the Coriell Institute for Medical Research in Camden, NJ. All control lines (n=14) are unrelated to the patient lines. Fibroblast cell lines were grown in DMEM (Gibco™#11995073), 10% FBS(Gibco™ #10-082-147), 1% Non-essential Amino Acid (100X, Gibco™, 11140-050) and 1% PenStrep (100X, Gibco™ #15140122) at 37℃ in 5% CO_2_ incubators. Cell lines were tested for mycoplasma on a monthly basis.

### RNAseq Library Preparation and Analysis

RNA was extracted from cells grown to 80-90% confluence using the PureLink RNA mini kit (Invitrogen #12183018A). Samples were processed for stranded total RNAseq libraries. The rRNA was depleted using the Qiagen QIAseq Fastselect-rRNA/Globin Kit (Qiagen #335377) followed by illumina TruSeq® Stranded Total RNA Library Prep Gold (illumina #20020599). Samples were multiplexed, and RNA sequencing was performed at the UCLA Technology Center for Genomics and Bioinformatics Core Facility at UCLA and sequenced on an Illumina NovaSeq for an average of 30 million reads per sample. Raw read quality, adaptor content, and duplication rates were assessed with FastQC. Raw reads were then aligned against the Gencode human genome version hg38 (GRCh38) version 31 using STAR 2.7.0e with default parameters (Dobin et al. 2013). Gene counts from raw reads were generated using featureCounts 1.6.5 from the Subread package. For each gene, we counted reads that uniquely mapped to the gene’s exons.

Differential expression was quantified using DESeq2 v1.34.0(Love et al. 2014). Genes with an adjusted p-value (Benjamini-Hochberg correction) less than 0.05 were considered as significantly differentially expressed (Wald’s test). LFC shrinkage of the DESeq object was done with apeglm (Zhu et al. 2019). A heatmap was created from normalized counts from the DEseq using R-package pheatmap (Kolde 2015), which shows the top differentially expressed (DE) genes based on adjusted p-values less than 0.05. Color on legend represents normalized counts. The top DE genes are listed in **Table S2**.

### ATACseq Library Preparation and Analysis

ATAC-seq was performed with 50,000 cells in both control and patient derived dermal fibroblast cell lines. ATAC-seq library generation was performed as described in (Corces et al. 2017). Samples were run on the Agilent DNA TapeStation to confirm tagmentation pattern, and the genomically barcoded libraries were multiplexed and run on the Hiseq3000 with paired end, 150bp libraries for a minimum of 30 million reads per sample.

ATACseq data was analyzed using an in-house bioinformatics pipeline (Lin et al. 2023). Briefly, the quality of reads were assessed using FastQC. Raw reads were then aligned to GENCODE Human genome version hg38 (GR38) version 31 using BWA-MEM (Li 2013). BAM files were then sorted, indexed, and filtered against chrX, chrY, and MT reads using *SAMtools*. *Picard* tools were then used to generate insert size histograms and remove duplicates from BAM files. Narrow peaks from each sample were called using MACS2 callpeak (Gaspar 2018); any peak that overlapped by at least one base was then merged using BEDtools (Quinlan and Hall 2010) *merge*. Reads overlapping merged peaks were counted using featureCounts (Liao et al. 2014). *DESeq2(Love et al. 2014)* was used to identify differentially open peaks between disease and control samples. Peaks with p-adj value (Benjamini-Hochberg) less than 0.05 were classified as significantly differentially open (Wald’s test), and fold changes were shrunk using approximate posterior estimation for GLM coefficients. LFC shrinkage of the DESeq object was done with apeglm (Zhu et al. 2019). Significant peaks were identified as promoter peaks if their distance from their respective closest gene was less than 1kb upstream or 2kb downstream relative to the gene transcription start site.

### ChIP-Seq Sample Processing and Analysis

The dermal fibroblast cells were grown to 80-90% confluency and then fixed using fresh media containing 1% formaldehyde. Cells were incubated in formaldehyde media for 10 minutes, and then the reaction was neutralized with 1.25M glycine. After this, the media was removed and cells were washed twice with ice cold PBS containing protease inhibitors. Post PBS wash, the cells were scraped and collected in a 1.5 ml tube. The cell pellets were resuspended in ChipLysis buffer and sonication was performed using Diagenode bioruptor at high for 45 cycles (30 sec on, 30 sec off). 5 ul of sonicated product was de-crosslinked and purified to check optimal lysis (200-600 bp).

The sonicated chromatin was pre-cleared with Antigen G dynabeads for 2 hours on a nutator at 4 ^0^ C. The pre-cleared chromatin was then incubated with 5 ul of H3K9ac (Cell signaling #9649), 2 ul of H3K23ac (AbCam #ab177275), and 4ul of IGg (Cell signaling #2729) at 4 ^0^C overnight. The next day, the antibody-chromatin complex was incubated with antigen G dynabeads for an additional 2 hours on a nutator, and then the beads were washed by a series of buffers and then de-crosslinked at 65 ^0^C overnight. Post de-crosslinking, the DNA was purified and processed for library preparation using illumina Chip library prep (Illumina#15034288). Chip libraries were sequenced on illumina Hiseq3000 using single read 75 bp read-length.

ChIP-Seq data was analyzed using an in-house bioinformatics pipeline similar to the ATAC-seq pipeline used above (Lin et al. 2023). In short, the quality of reads were assessed using FastQC. Raw reads were then aligned to GENCODE Human genome version hg38 (GR38) version 31 using BWA-MEM (Li 2013). BAM files were then sorted, indexed, and filtered against chrX, chrY, and MT reads using *SAMtools*. *Picard* tools was used to remove duplicates from BAM files. Narrow peaks from each sample were called using MACS2 callpeak (Gaspar 2018) by using the control sample BAM to account for nonspecific antibody parameter *-c*; based on resulting peaks called, any peak that overlapped by at least one base was then merged using BEDtools (Quinlan and Hall 2010) *merge*. Reads overlapping merged peaks were counted using featureCounts (Liao et al. 2014). Samples with over 15 million reads were kept.

*DESeq2 (Love et al. 2014)* was used to identify differentially acetylated peaks between disease and control samples. Peaks with p-adj value (Benjamini-Hochberg) less than 0.05 were classified as significantly differentially captured (Wald’s test), and fold changes were shrunk using approximate posterior estimation for GLM coefficients. LFC shrinkage of the DESeq object was done with apeglm (Zhu et al. 2019).

### DNA methylation Library Preparation and Analysis

DNA methylation generated from ARTHS and control fibroblasts were processed as described in (Awamleh et al. 2022; Bondhus et al. 2022). Briefly, DNA was extracted from both control and patient derived dermal fibroblast cell lines. The extracted DNA was bisulfite converted and run on the Illumina MethylationEPIC Array as previously described (Mansell et al. 2019) at the UCLA Neuroscience Genomics Core. The MINFI package (Aryee et al. 2014) was used to perform QC on the resulting idat files. Probes overlapping SNPs and those on the sex chromosomes were filtered out. After QC and filtering, 832,159 measured CpGs remained of the 865,919 measured CpGs included on the MethylationEpic Array. Background correction (Triche et al. 2013) and functional normalization (Fortin et al. 2014) were used for preprocessing and normalization of individual probes.

### Gene Ontology Enrichment

Gene ontology over-enrichment tests were completed using g:Profiler (Raudvere et al. 2019) by submitting the either the Differentially expressed genes or the closest genes to significantly differentially open peaks against all genes from the Gencode hg38 annotation, version 31. Gene ontologies were classified as significantly enriched when p-adj (Benjamini-Hochberg) was less than 0.05 (hypergeometric test).

### Cytoplasmic and Nuclear Protein Extraction

The cells were harvested using trypsin-EDTA and centrifuged at 500 X g for 5 minutes, and the cell pellet was washed with PBS. The dry cell pellet was then processed for nuclear and cytoplasmic protein extraction as per manufacturer’s protocol (Thermo Scientific™NE-PER™ Nuclear and Cytoplasmic Extraction Reagents, #78833). For western blotting assay, 10 ug of the nuclear extract protein were loaded on the 4–15% Criterion™ TGX Stain-Free™ Protein Gel (Bio Rad #5678083). The protein was transferred to the nitrocellulose (Bio Rad # 1704271) using a turbo-transfer system, blocked with 1X TBST with 5% non-fat milk for an hour at 4℃, and then incubated with Anti-KAT6A / MOZ antibody (Active Motif, at 1:500) overnight. The blots were then washed and detected with IRDye® 800CW secondary antibodies (LI-COR #926-32210 and # 926-32213). For control antibody, the blots were incubated in and hFAB™ Rhodamine Anti-Actin (BioRad #12004163 at 1:2000 dilution). All experiments were performed in triplicate.

### Histone Extraction

For histone extraction, the cells were washed twice with ice-cold dPBS (Gibco #14190144) containing 0.5nM sodium butyrate. 750μL of prepared lysis buffer (10mM HEPES, 1.5 mM MgCl2, 10mM KCl, 0.34M Sucrose, and 10% glucose) with fresh 1x protease (Thermofisher, 78442) and 1mM of DTT (Thermofisher, A39255) were added to each plate. Cells were scraped and transferred to an eppendorf tube, and 1% Triton-X was added to a final concentration of 0.1%. Tubes were incubated on ice for 8 minutes followed by centrifugation for 5 minutes at 4°C and 4500xg. After aspirating the supernatant, the nuclei were lysed by adding 750μL of lysis buffer, and then spun again at 4°C at 4500 x g for 2 minutes. The supernatant was aspirated, 50μL of extraction solution (50μL 10% glycerol, 6uL concentrated H2SO4, 1μL BME, and 443μL water) was added to the pellet, the tubes were incubated on ice for 10 minutes, and then centrifuged at top speed for 10 minutes at 4°C. 50μL of the supernatant was transferred to a new tube, and 12.5μL of 100% TCA was added. After being vortexed briefly, the tubes were centrifuged at 4°C for 10 minutes at top speed. The supernatant was aspirated, and 1mL of cold ethanol was added. The tubes were vortexed, and then incubated at −80°C for 10 min before being centrifuged again at 4°C for 10 min at 16,000 x g. The ethanol was aspirated and the pellets were allowed to air dry for at least 20 minutes. Histone pellets were dissolved in 100μL water. For quantification of histone acetylation levels, 250ng - 1ug of histones were loaded on the gel and were detected for H3K9ac (Cell signaling#9649, 1:1000 dilution) H3K23ac (Cell signaling#8848, 1:1000 dilution) and H3core(Cell signaling#3638,1:1000 dilution)

### Histone Extraction for Mass Spectrometry

Fibroblasts isolated from unaffected controls (n=3) and ARTHS patients (n=3) were grown as previously described until cells were approximately 80-90% confluent. Cells were then harvested, washed once with DPBS, pelleted, snap-frozen, and stored at −80°C for subsequent histone extraction and mass spec analysis. Histones were extracted from snap-frozen pellets using a previously published protocol (Sidoli et al. 2016). Briefly, the cells were thawed and lysed on ice using a nuclear isolation buffer (NIB) containing NP-40. After lysis the cells were centrifuged and washed in NIB without NP-40 to obtain pure nuclear extracts. After this the nuclei were resuspended in 0.2M H_2_SO_4_ and incubated on constant rotation a 4^0^C for 2-4 hours. Then the samples were centrifuged at 3400Xg for 5 minutes and supernatants were collected in fresh tubes. The Histones were precipitated using chilled 100% TCA and incubated for either 1 hour/overnight. The sample tubes were centrifuged and the nuclei pellets were washed with ice cold acetone and dried and dissolved in nuclease free water. Once the histones were quantified, about 20 ug of histones were dissolved in 50mM NH_4_HCO_3_ pH 8.0 to a final concentration of 1ug/ul. Then fresh propionylation reagent was added to the sample in 1:4 v/v ratio, NH_4_OH was quickly added in 1:2 v/v ratio to keep maintaining the pH 8.0. The samples were vortexed and incubated for 15 minutes at room temperature. The samples were then completely dried down using a vacuum evaporator and resuspended in 50mM NH_4_HCO_3_ pH 8.0 to achieve a concentration of 1ug/ul of protein. The propionylation procedure was performed again for a total of two rounds The samples were digested using trypsin (1:50 ratio wt/wt) and incubated for 6-8 hours at 37℃. The Digestion was stopped by storing the samples at −80℃. The samples were dried down to 10-20ul using vacuum and then resuspended in 50mM NH_4_HCO_3_ to make up the volume to 20ul. After this the samples undergo another two rounds of propionylation as mentioned above. The samples were desalted by C18 stage tips, dried down, and resuspended in 0.1% formic acid to a final concentration of 0.5 ug/ul.. Once the samples are ready they are analyzed using LC-MS.

## Funding

This work was supported by the following funding sources awarded to V.A.A.: NIH DP5OD024579, the UCLA Eli and Edythe Broad Center of Regenerative Medicine and Stem Cell Research Rose Hills Foundation Innovator Award 2022-2023, the funding sources to B.G. NIH NS111997 and NIH HD106051, and the following funding sources awarded to A.A.N.: the Graduate Dean’s Scholar Award Fellowship (2019-2021), the Broad Stem Cell Research Center Training Fellowship (2021-2022) and the Eugene V. Cota-Robles Fellowship (2019-2023)

### Contributions

M.S. and V.A.A. conceptualized and coordinated this study. M.S. performed experiments and designed the various sequencing approaches. A.A.N., F.V. and B.G. performed and analyzed proteomics experiments. J.L. and G.E. performed western experiments. S.S. and A.W. analzyed RNAseq, ATAC-seq, and ChIPseq studies. L.B. and S.A. analyzed DNA methylation data and Motif binding analysis. All authors contributed to the writing and editing of the paper.

## Acknowledgements

This work would not be possible without the support and generous participation of the *KAT6A* foundation and all the patients and their families for their donation of samples over the past years. We also thank Stanley Nelson, Emilie Douine and Elisabeth McGee of the California Center for Rare Disease, who maintained and coordinated our human research studies over the years. Finally, we acknowledge the use of the Pathology Research Portal for their support in processing samples that arrived from all over the country.

## Data Sources

DNA methylation study is available under GEO record GSE210484. RNAseq, ATAC-Seq and ChIPseq data from this study is uploaded into GEO study accession number GSE237023.

