## supplemental figures for "*KAT6A* mutations in Arboleda-Tham syndrome drive epigenetic regulation of posterior *HOXC* cluster"

**AFFILIATIONS**
1 Department of Pathology & Laboratory Medicine, David Geffen School of Medicine, UCLA, Los Angeles, CA, USA
2 Department of Human Genetics, David Geffen School of Medicine, UCLA, Los Angeles, CA, USA
3 Department of Computational Medicine, David Geffen School of Medicine, UCLA, Los Angeles, CA, USA
4 Interdepartmental BioInformatics Program, UCLA
5 Department of Biochemistry and Molecular Biophysics, Washington University in St. Louis

**Corresponding Author:**
Valerie A. Arboleda
Associate Professor
Departments of Pathology & Laboratory Medicine, Human Genetics and Computational Medicine
David Geffen School of Medicine UCLA
615 Charles E. Young Drive South
Los Angeles, CA 90095


Supplementary figure 1:

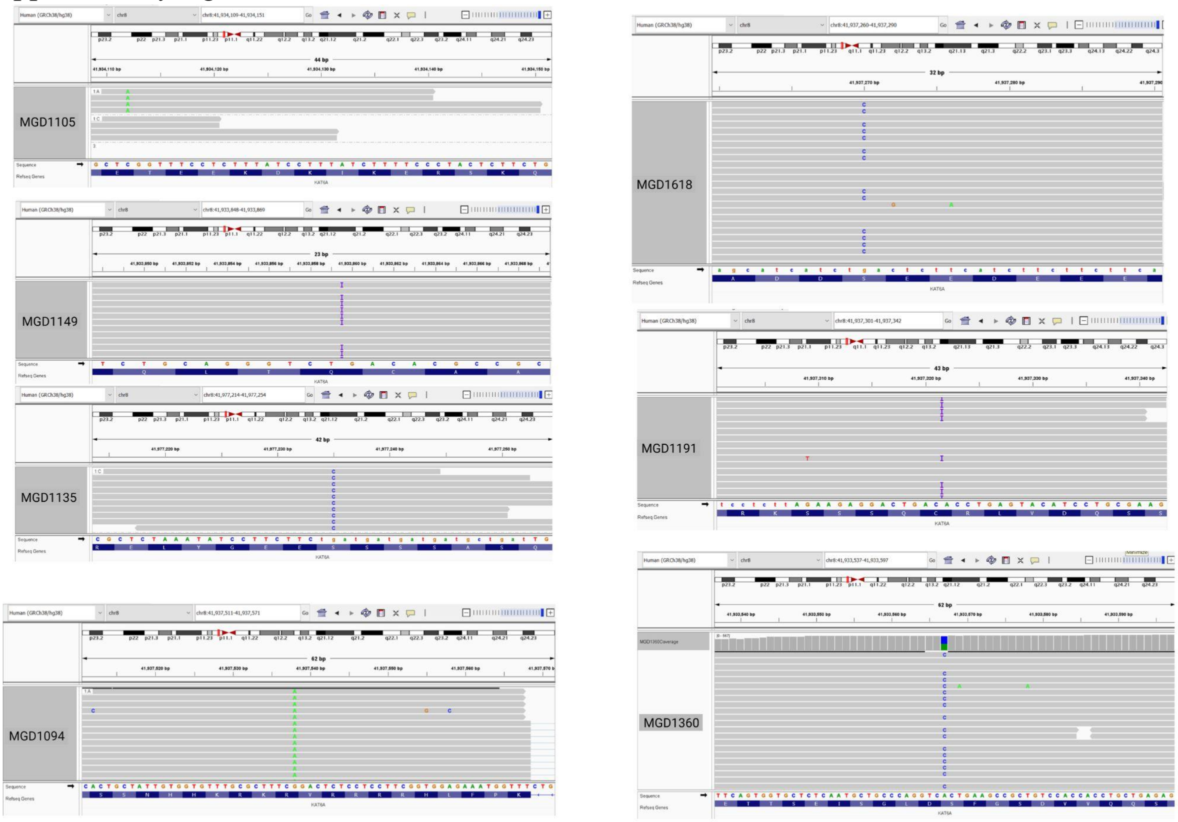


**Supplementary Figure 1: Genome track files confirming the ARTHS patient mutations.** All 7 ARTHS patient RNAseq data was uploaded onto the IGV browser to confirm the mutations.

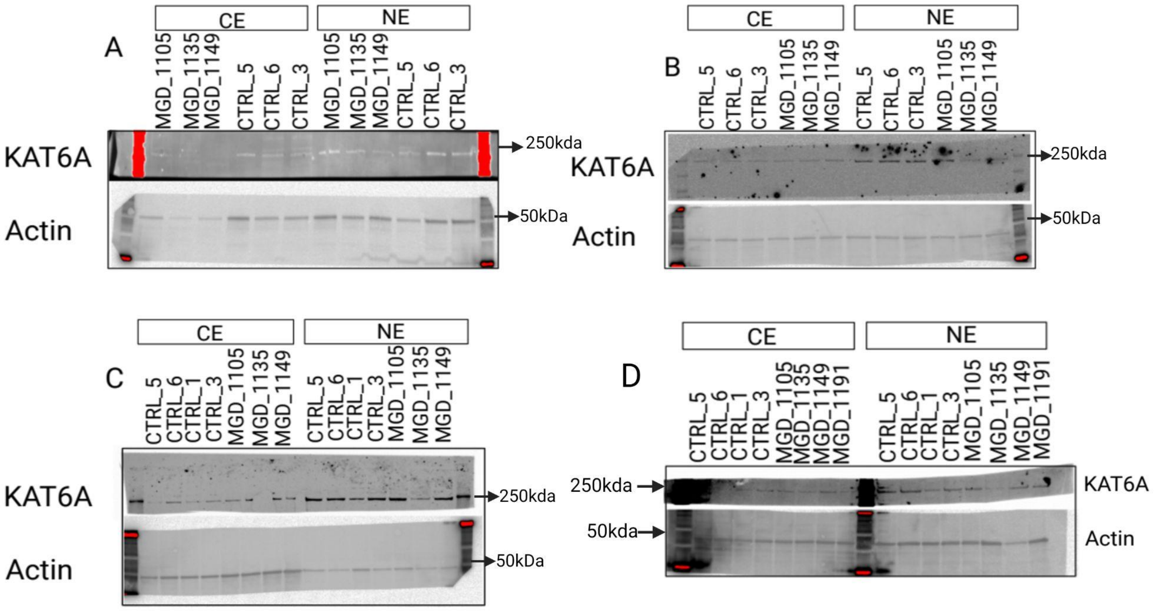


**Supplementary Figure 2**: Western blots of KAT6A protein levels in cytoplasmic and nuclear fractions of cell pellets do not show significant changes in the ARTHS fibroblast samples as compared to the controls. A, B, C, and D represent 4 replicates used in quantification. KAT6A (225 kDa), Actin (42kDa).

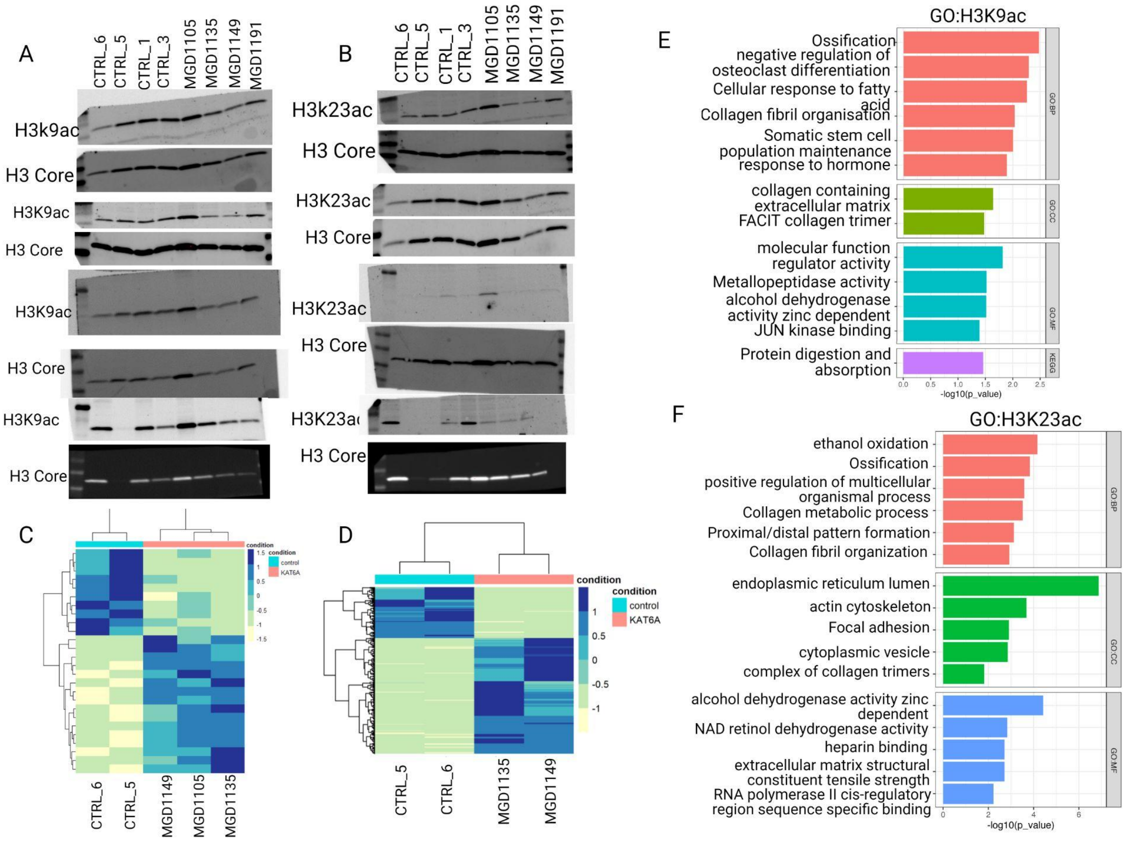


**Supplementary Figure 3: ARTHS mutations do not change global acetylation levels at H3K9 or H3K23.** A) Western blot images for the H3k9ac mark across 4 independent blots in a subset of ARTHS and control samples. These are quantified in Figure 4B. B) Western blot images for the H3K23ac mark across a subset of ARTHS and control samples. These are quantified in Figure 4C. Heatmap depicting differentially acetylated peaks from ChIP-seq data at the C) H3K9ac mark and D) H3K23ac mark. E) GO terms associated with differentially acetylated genes in the H3K9ac ChIPseq dataset F) GO terms associated with differentially acetylated genes in the H3K23ac ChIPseq dataset

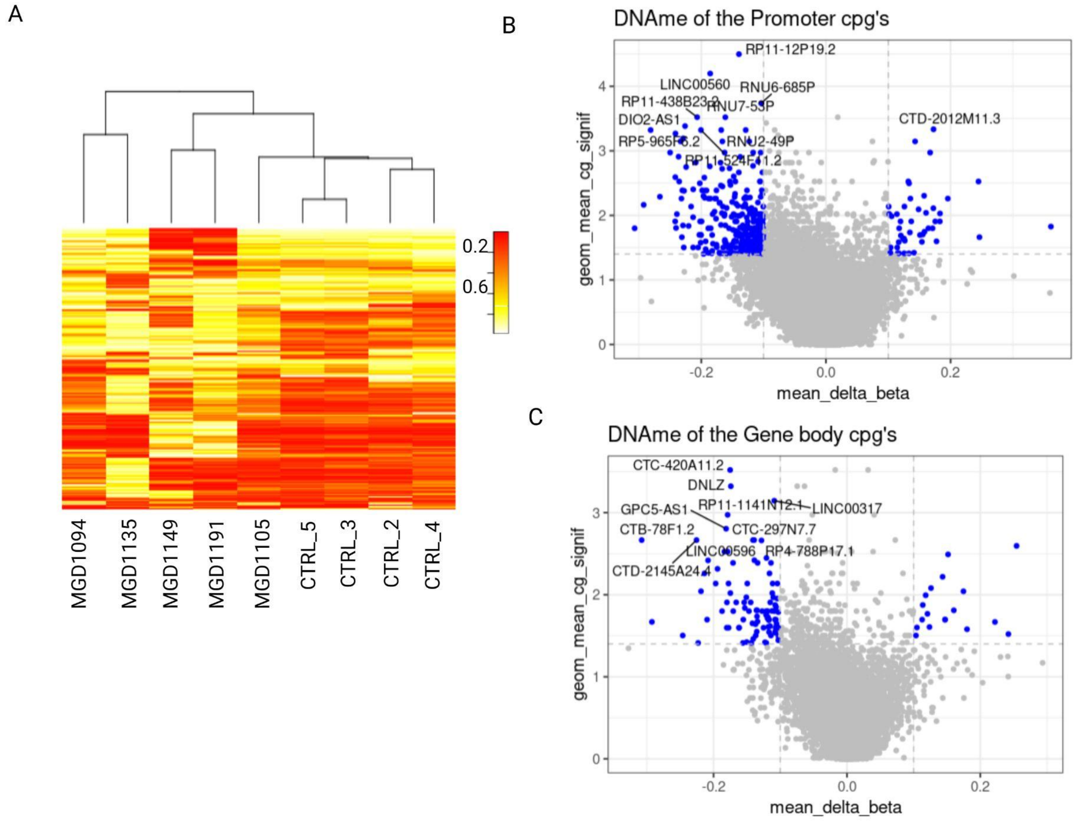


**Supplementary Figure 4:** CpG methylation profiling of ARTHS samples: A)Heatmap showing sample clustering and the levels of DNA methylation across ARTHS and control fibroblasts for the top 1000 most variable CpGs. B) Volcano plot showing promoter CpGs that are differentially methylated between ARTHS and control fibroblasts. Significantly different promoter CpGs are highlighted in blue. C) Volcano plot showing gene body CpGs that are differentially methylated between ARTHS and control fibroblasts. Significantly different genebody CpGs are highlighted in blue.

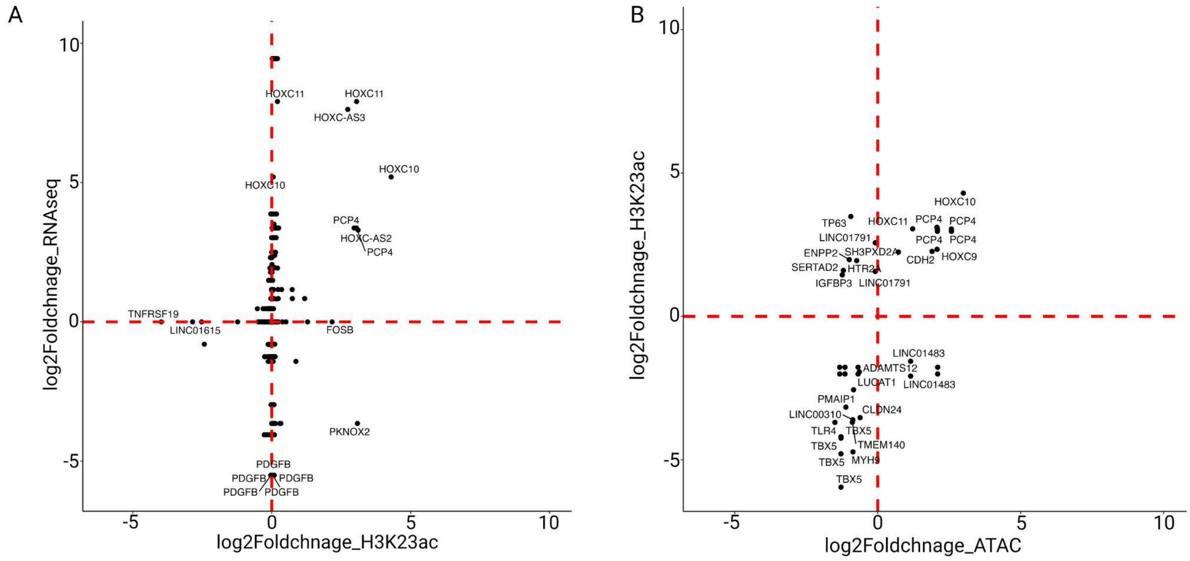


Supplementary Figure 5: Multi Omic data integration suggests that ARTHS KAT6A mutations regulate the expression of posterior HOXC cluster genes. A) Correlation between H3K23ac ChIPseq and RNAseq data. It highlights the expression change and H3k23 acetylation being changed on *HOXC* genes. B) Correlation betweenH3k23ac ChIPseq and ATACseq data. It highlights increased H3k23 acetylation and increased chromatin accessibility over the posterior *HOXC* genes.
